# Long-term *ex ovo* culture of *Caenorhabditis elegans* embryos

**DOI:** 10.1101/2025.06.26.661779

**Authors:** Clover Ann Stubbert, Cherry Soe, Pavak Kirit Shah

## Abstract

While the genetic tractability, transparency and invariant development of the *Caenorhabditis elegans* embryo have led to its broad adoption as a model system for the study of cell and developmental biology, its impermeable eggshell has complicated the use of small-molecule reagents during embryogenesis. Existing genetic approaches for rendering the embryo permeable to acute small molecule treatment have increased the accessibility of early embryogenesis to pharmacological manipulation but compromise long-term viability, preventing their use in studies of later developmental processes or post-exposure physiology. Here, we describe the use of an optimized enzymatic eggshell digestion protocol coupled with a minimal, serum-free culture medium that supports the survival and normal development of *ex ovo* embryos through larval maturation and adulthood. We show that this approach renders embryos permeable to a wide range of small molecules, enabling precise temporal manipulation of developmental processes previously inaccessible through conventional genetic methods. We demonstrate the utility of this technique through the pharmacological modulation of cytoskeletal components including microtubules and actin, as well as the minus-end-directed microtubule motor protein dynein, highlighting applications for the study of cell division, morphogenesis, and neuronal development, especially at later stages of embryogenesis. As a proof of concept, we use acutely timed dynein inhibition to show that cytoplasmic dynein is required to transport the centriole into the dendrite of embryonically born sensory neurons. This approach expands the experimental toolkit available for labeling and manipulating developmental processes in *C. elegans*.

## Introduction

The nematode *Caenorhabditis elegans* is a powerful model system for the study of genetics, development, and cell biology owing to its genetic tractability, transparency across life stages, and invariant development. In contrast to the amenability of the *C. elegans* embryo to live imaging^1,2^, advances in the perturbation of developmental processes have been primarily limited to genetic^3,4^ and optical approaches^5–8^ by the impermeability of its eggshell, consisting of multiple proteoglycan and chitin layers^9^. This renders *C. elegans* embryos impermeable to most small molecules.

A genetic screen previously identified multiple genes (*perm-1, perm-2*) required for the eggshell permeability barrier^10^. Knock-down of these genes by RNAi causes the generation of embryos whose eggshells are permeable to small molecules, which has enabled a wide range of pharmacological manipulations of the early embryo^11–15^. This approach has some drawbacks, as it results in small brood sizes and compromised viability with embryos typically failing to develop beyond gastrulation^10,16^, making it unusable for the labeling and manipulation of later developmental stages. Physical methods of permeabilizing the eggshell (laser ablation^17,18^, acute coverslip pressure^19,20^, and microinjection^21^) also have been used to introduce small molecules into *C. elegans* embryos but are limited in throughput and introduce sources of variability such as the magnitude of applied pressure and the targeting of the laser microbeam.

We reasoned that an approach based on cell culture techniques may be able to support the long-term culture of fully eggshell-removed, yet intact embryos. Indeed we have found that optimized eggshell digestion protocols, when combined with a simplified serum-free culture medium, supports the survival and growth of embryos through larval maturation, producing healthy individuals that are able to be transferred to standard culture plates and raised to fertile adulthood. Since others have previously shown that chitinase digestion is sufficient to render embryos permeable to small molecules^22^, we tested whether these cultured embryos are compatible with manipulation by many common small molecules (eg., toxins, and selective inhibitors) at developmental stages that are not accessible by *perm-1* and *perm-2* RNAi. We demonstrate that these treatments can be used to manipulate a wide range of biological processes and structures within the *C. elegans* embryo including the microtubule and actin cytoskeleton and the activity of the minus end-directed microtubule motor protein dynein. This work provides a scalable approach that expands the toolkit available for the labeling and manipulation of the *C. elegans* embryo for studies of development and cell biology. As a proof of concept that *ex ovo* culture opens developmental events that are inaccessible to conventional genetics, we used acute, temporally precise dynein inhibition to test whether dynein activity is required to transport the centrosome into the dendrite of embryonically born sensory neurons, complementing prior genetic analysis of the postembryonic PQR neuron^23^.

## Results

### *Ex ovo* culture of *C. elegans* embryos

Genetic perturbations that render the eggshell permeable to small molecules also reduce viability^10^, making these approaches incompatible with studies of developmental processes that occur after gastrulation such as tissue morphogenesis and neuronal development. We hypothesized that media optimized for nematode embryonic cell culture may support embryo viability in the absence of the eggshell permeability barrier. We use a simplified version of a culture medium originally developed for maintaining *C. elegans* neuronal and muscle cells^24^, omitting the serum the original protocol used so as not to provide the intact embryo with exogenous growth factors. This minimal medium is prepared from commercially available Leibovitz’ L-15 medium supplemented with 7.7 g sucrose / 500 mL and 1× penicillin and streptomycin (can be omitted for short-term culture).

After digestion by Yatalase, the eggshell is clearly missing when observed under high magnification DIC microscopy. Given the handling time needed to dissect early-stage embryos, transfer into Yatalase and recover after digestion, we have not succeeded at reliably preparing the single cell zygote in this manner. 2-cell embryos can be reliably isolated, digested and survive in L-15 + sucrose at rates comparable to later stage embryos (29/31). These early embryos digest quickly with 30% embryos having no visible eggshell within 5 minutes. To obtain the quickest digestion possible, roughly 30 embryos were added to the yatalase solution and immediately imaged under DIC to determine the timing of digestion. Pre-bean stage embryos show no visible remaining layers under DIC, but at comma stage, a flexible membrane becomes visible surrounding the embryo, which may be the peri-embryonic layer observed in electron micrographs of the eggshell^9^. Whether this membrane is formed at these stages or is simply too tightly bound to the embryo surface at earlier stages to be visible by DIC is unclear. These embryos hatch at similar times to undigested *C.elegans* embryos. Upon completing embryonic development, L1 larvae “hatch” from this soft membrane as normal (Fig 1A). We observe that >80% of early-stage embryos (younger than the bean stage and thus lack an epidermis at the time of eggshell digestion) survive and hatch as L1 (40/50) (Fig 1G). Many of the embryos that fail to hatch show signs of mechanical trauma, likely suffered during the micropipette transfer process. We recovered a portion of the L1 larvae onto standard NGM plates seeded with OP50 *E. coli* and observed that >90% of those survived to become fertile adults (28/30) (Fig 1G). Late-stage embryos are approximately as successful in reaching L1 (35/50) and becoming gravid adults (29/31), demonstrating that this minimal culture medium is capable of maintaining embryo viability after eggshell removal at any stage of embryo development starting at the 2-cell stage and that animals treated in this fashion are fully capable of surviving and maturing to adulthood. The few larvae that are lost were not found, suggesting they were likely lost under the medium or up the petri dish side walls. Unsupplemented L-15 and standard egg buffer^25^ (commonly used medias for *C.elegans* cell and embryo culture) yield a much lower rate of survival to L1 (46% (20/46, 27/53, 20/48) and 48% (9/17, 26/52, 24/56) respectively) with arrested embryos showing signs of severe osmotic stress such as swelling and bursting (Fig 1B-C).

**Figure 1.**
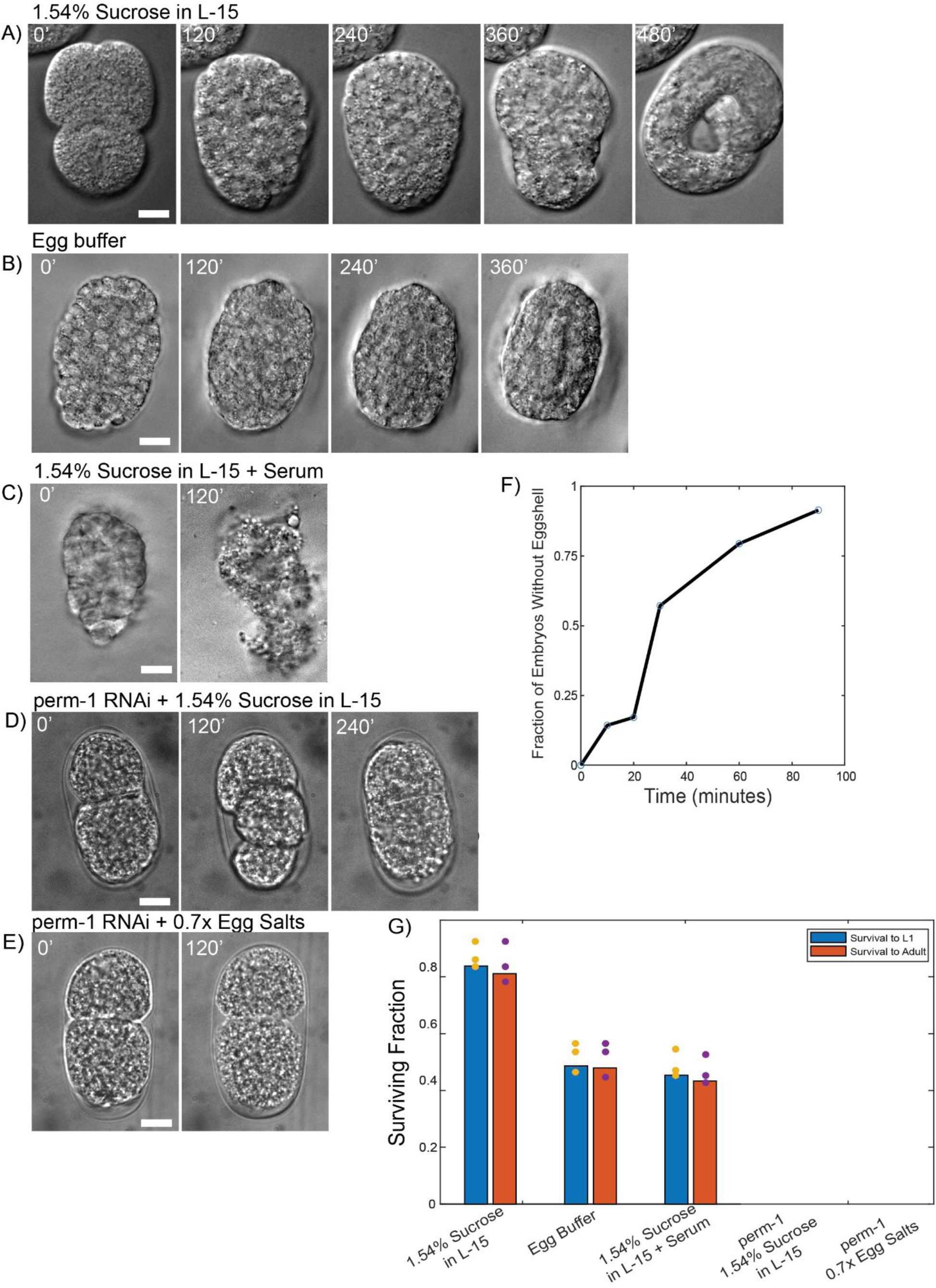
A) *Ex ovo* embryos develop normally in 1.54% sucrose in L-15 (10 μm scale bar). B) *ex ovo* embryos show signs of osmotic stress when left in standard egg buffer and arrest (10 μm scale bar). C) *ex ovo* embryos in 1.54% sucrose L-15 medium with serum quickly arrest and undergo apoptosis (10 μm scale bar). D) perm-1 embryos cultured in 1.54% sucrose in L-15 (10 μm scale bar). E) perm-1 RNAi embryos in 0.7X Egg Salts (10 μm scale bar) F) Line plot showing the increase in digestion over time. G) Survival rates of *ex ovo* embryos cultured overnight in 1.54% sucrose in L-15, egg buffer, and serum supplemented L-15 medium and perm-1 embryos in 1.54% sucrose in L-15 and in 0.7x Egg Salts. Blue bars show survival to L1 while orange bars show survival to fertile adulthood.

We next optimized Yatalase digestion duration and yield. By observing embryos in the chitinase solution periodically under DIC microscopy, we tracked the progression of eggshell digestion over time by observing when the eggshell is no longer visible. When digested in batches of 50 embryos in 12.5 mg/mL Yatalase, just over half of the embryos are fully digested within 30 minutes with an increase to roughly 80% of embryos digested within 90 minutes (Fig 1F). The amount of time that embryos are rocked in Yatalase to digest can be shortened or lengthened depending on the desired yield of fully digested embryos. Reducing the number of embryos to 20 per batch accelerated digestion, with 80% showing no visible eggshell within 60 minutes. It is likely that either inactivation of the Yatalase or the increasing concentration of degraded chitin may be limiting the reaction. We tested this by replacing 20% of the chitinase suspension with fresh enzyme solution after 30 minutes of incubation and observed a marked increase in the ratio of digested embryos in large embryo preps (n > 100 embryos) 20 minutes following the addition of the enzyme (removal of 10 µL and readdition of 10 µL of enzyme solution for total volume of 50 µL). This strategy reduces the survival rate of embryos to 58% for early-stage embryos (29/50) and 54% for late-stage embryos (27/50 embryos) but of those that survived to L1 roughly 90% of both early and late-stage embryos became gravid adults (E=27/29, L=23/27). This drop in survival rates can likely be attributed to the extra handling of the embryos and additional shear stresses during liquid exchange, as even partial eggshell digestion renders embryos extremely fragile.

Knockdown of *perm-1* using RNAi is commonly used to permeabilize early-stage embryos for small molecule treatment. Using the *perm-1* RNAi, we assayed survival in the culture conditions reported (0.7% egg salts) and saw that all (0/50) embryos tested failed to survive past the two-cell stage (Fig 1D). Culturing L15 + sucrose did not improve survival in *perm-1* RNAi with all embryos similarly failing to develop (0/50) (Fig 1E).

### Manipulation of microtubule stability in *ex ovo* embryos

The microtubule cytoskeleton plays a critical structural and functional role in a wide range of fundamental cellular processes from mitosis^26^ to morphogenesis^27^. Genetic and biochemical studies of *C. elegans* have identified a range of functional regulators of microtubules^28,29^. The essential role of microtubule regulation creates challenges for the study of the cytoskeleton at later developmental stages. The incomplete permeability barrier of the early single cell embryo and laser microbeam based permeabilization have previously been used to deliver microtubule toxins and drugs targeting microtubule regulators^10,30^. We reasoned that this approach would also make it possible to utilize microtubule toxins to manipulate the cytoskeleton at a wider range of developmental stages, without laser microbeam or coverslip pressure-assisted permeabilization.

Taxol functions by stabilizing microtubule filaments, preventing them from depolymerizing and stopping cell divisions causing mammalian cells to undergo apoptosis^31^. It has previously been used to study spindle axis establishment of the first cell division of the embryo in *C. elegans* and used as a control when testing new destabilizing treatments^30,32^. We tested a range of taxol doses in *ex ovo* embryos. Starting with a dose that had previously been used with *perm-1* RNAi treated embryos (100 nM) (n = 13 embryos), all cells immediately ceased visible motion and did not divide further with the same effect being seen at 50 nM (n = 5 embryos). The addition of 100 nM taxol to embryos with intact eggshells caused no visible phenotype leading us to believe it is not able to penetrate the eggshell. Lower doses (10 nM) slowed the onset of complete stabilization treatment and allowed for the execution of a single post-treatment round of cell division to be completed before ceasing motion and further division (n = 5 cells across 5 embryos) (Fig 2B). Using even lower doses (5 nM) (n = 7 embryos), we observed an increase in the amount of tubulin associated with the mitotic spindles in early-stage embryos, with the spindles pulled outward to the cell cortex (Fig 2A). At the lowest dose we tested (1 nM), cell divisions progressively slowed over the course of an hour until, regardless of embryo stage, all embryonic cell divisions stopped (n = 5 embryos). In all cases, 100% of embryos lacking an eggshell based on DIC observation showed these responses to taxol treatment.

**Figure 2.**
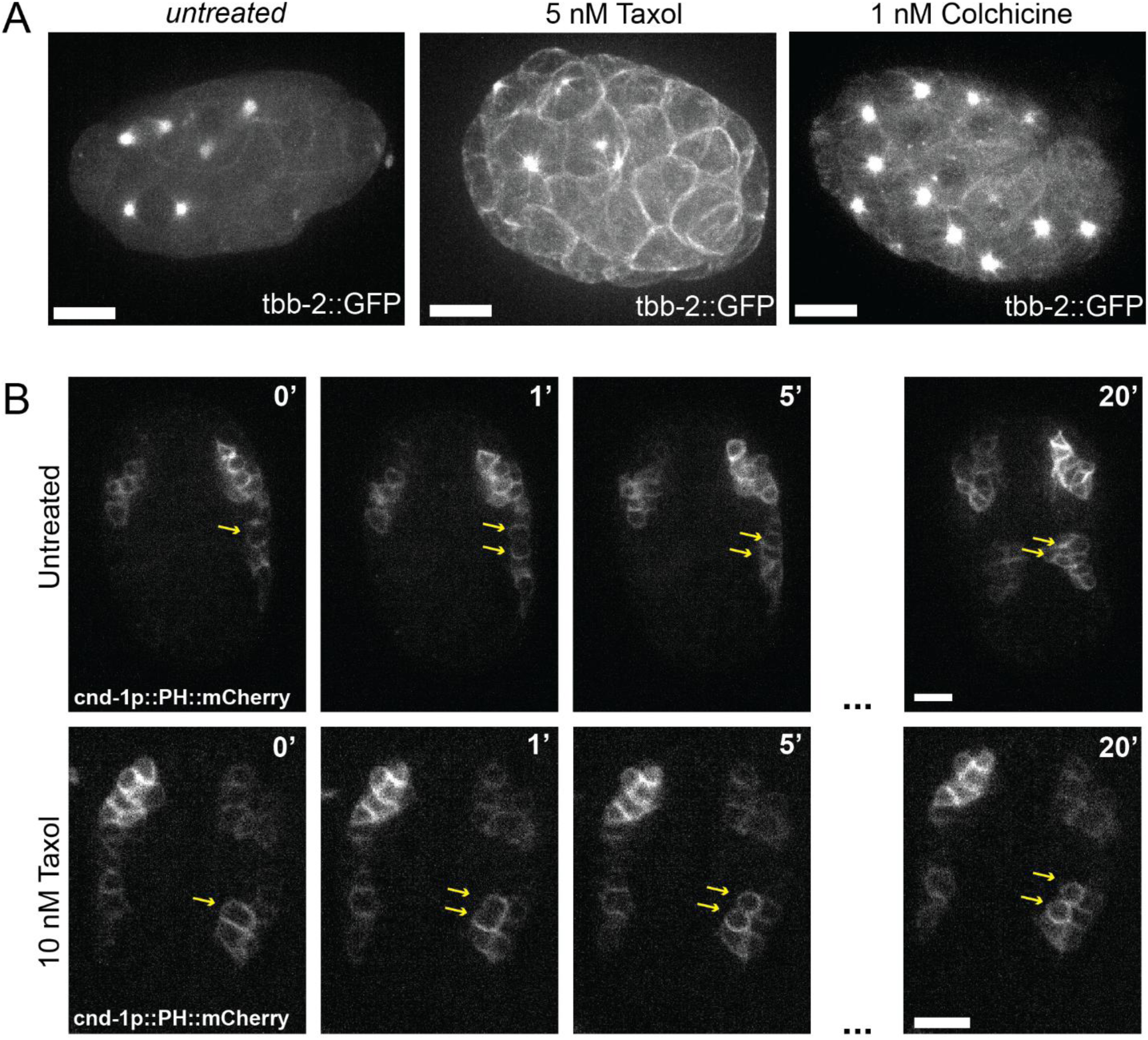
A) Maximum projection of tbb-2::GFP early-stage *ex ovo* embryo in the absence of toxin (left). Maximum projection of tbb-2::GFP early-stage embryo treated with 5 nM Taxol (middle). Maximum projection of tbb-2::GFP early-stage embryo treated with 1 nM of colchicine (right). Image intensity scaling is identical for all three images. B) Maximum projections of cnd-1p::PH::mCherry untreated embryos and embryos treated with 10 nM taxol at the time of treatment, one minute post treatment, two minutes post treatment, five minutes post treatment and twenty minutes post treatment. Yellow arrow heads pointing to cell division post treatment.

Colchicine is a microtubule depolymerizer at high concentrations and a stabilizer at low concentrations^33^. In adult worms, colchicine has been used to disrupt dendritic microtubules in sensory neurons to determine their function in sensitivity to stimuli^34^. At 1 nM, Colchicine mimicked the effects of Taxol treatment, causing a progressive accumulation of β-tubulin at mitotic spindles and a slowing of cell divisions leading to cessation of intracellular motion and division within 30 minutes (n = 17 embryos) (Fig 2A).

### Manipulation of actin polymerization in *ex ovo* embryos

The actin cytoskeleton regulates many active processes in cell shape change and migration. The regulation of actin polymerization and branching generates mechanical anisotropies that direct a wide range of developmental events and has been extensively studied in the context of hypodermal enclosure in *C. elegans*^35,36^. Following gastrulation, hypodermal cells migrate towards the midline to enclose the embryo by extending actin-rich protrusions along the leading edge, which is unaffected in *ex ovo* embryos (protrusions marked by arrowheads) (n = 10 embryos) (Fig 3A). Both the microtubule cytoskeleton and the assembly of contractile actin-myosin filaments are needed for proper enclosure. Embryos that fail to properly enclose typically rupture^36,37^.

**Figure 3.**
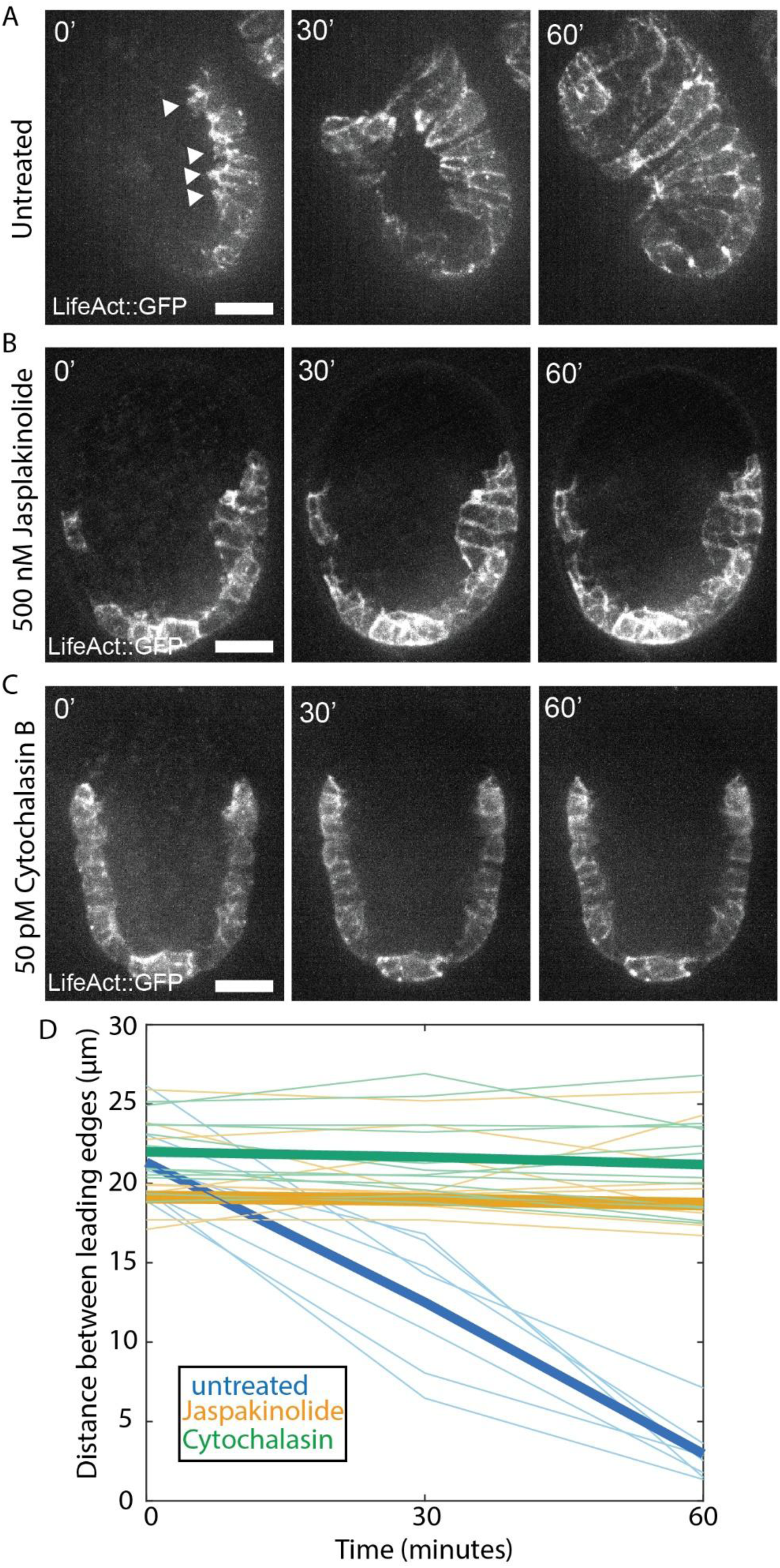
A) Maximum projection of untreated embryos in the absence of drug. B) Maximum projection of *ex ovo* embryos following treatment of 500 nM Jasplakinolide. C) Maximum projection of *ex ovo* embryos following treatment of 0.05 nM Cytochalasin B. D) Line plot showing the distance between leading edges of enclosing embryos over time in untreated (blue), Jasplakinolide (orange) and Cytochalasin B (green) treated embryos.

Jasplakinolide acts by binding to and stabilizing f-actin. It has been used to implicate F-actin contractile rings in the differentiation of spermatid^38^. At high doses, matching its prior use in adult *C. elegans*, (500 nM), Jasplakinolide treatment in embryos at the beginning of ventral enclosure blocked ventral migration and abolished ventral-oriented protrusions in the hypodermis of all treated embryos (Fig 3B) (n = 7 embryos)^39^. Treating embryos with intact eggshells had no visible effect (n = 9 embryos). Lower doses of Jasplakinolide (10 nM and 1 nM) also resulted in a complete block of ventral enclosure (n = 12 embryos and n = 8 embryos respectively).

Cytochalasin B is an inhibitor of actin polymerization by binding to F-actin and physically blocking filament extension. The addition of 200 nM Cytochalasin B, the dose used previously in laser microbeam permeabilized embryos, to embryos with an intact eggshell at the beginning of ventral enclosure results in no visible phenotype (n = 20 embryos)^19^. The lowest dose we tested (50 pM) also resulted in a fully penetrant failure of ventral enclosure (n = 8 embryos) (Fig 3C/D).

### Dynein is required for the dendritic transport of the embryonic centrosome

Centrosomes must maintain integrity under the high levels of cortical stress applied during mitosis as it forms the mitotic spindle^40,41^. Cytoplasmic dynein plays an important role in shuttling proteins from the cytoplasm to the pericentriolar material to maintain centrosome integrity and to mediate centrosome duplication^42,43^. Previous studies have shown that inhibiting cytoplasmic dynein will cause cells to fail to divide and centrosomes will lose structural integrity leading to an increase in the size of the pericentriolar material^42,44,45^. Prior work across cell culture, mammalian neurons, *Drosophila* neuroblasts and *C.elegans* sensory neurons have demonstrated that the minus end directed microtubule motor protein dynein is necessary for transport of the centrosome to the site of ciliogenesis. In *C. elegans,* a subset of sensory neurons execute a terminal division after the neuroblast extends a dendrite, which was recently confirmed in electron-microscopy (EM) reconstructions^23,46,47^. Like mammalian olfactory sensory neurons, whose centrioles migrate long distances to the apical surface for ciliogenesis^48^, these cells must transport the centrosome along the pre-existing dendrite to the site of ciliogenesis^23^. EM reconstructions further show that not all dendrites at a given stage contain a centrosome^47^, indicating a defined window of centrosome transport after the terminal division. In these cells, the centrosome translocates from the cell body to the sensory ending following this division (Fig 4L). Prior work established a role for dynein in centriole transport during dendrite morphogenesis, relying on bulk heat shock-induced targeted mutagenesis of the dynein heavy chain *dhc-1*, and then assessing larval animals for basal body positioning defects^23^. We asked whether dynein activity is similarly required for long-range transport of the centrosome in embryonically born sensory neurons using *ex ovo* embryos and a small molecule inhibitor of dynein.

**Figure 4.**
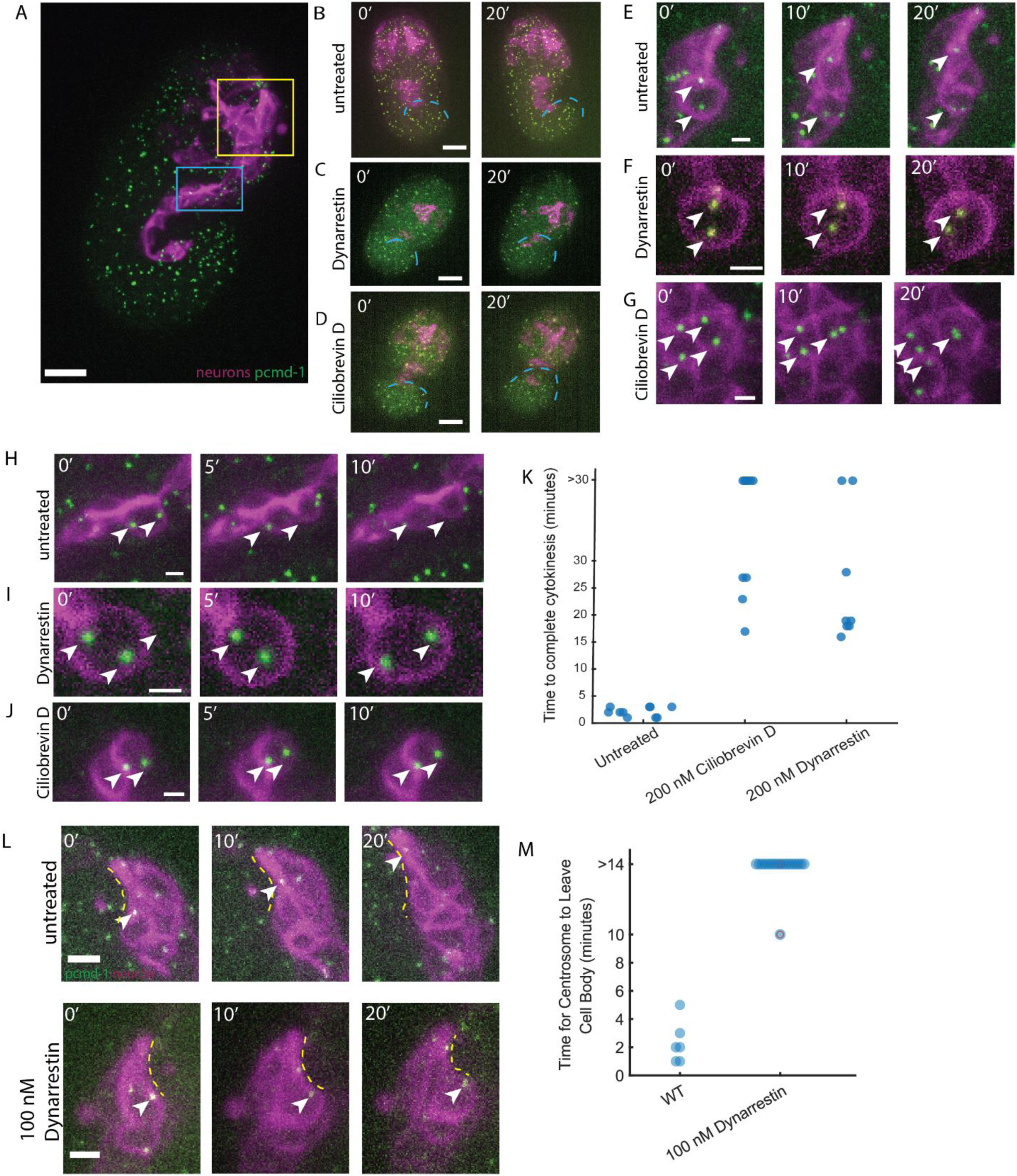
A) maximum projection of PCMD-1::GFP embryo. Yellow box showing head neurons. Blue box showing ventral nerve cord (VNC) Scale bar is 10 μm. B) Max projection of untreated embryo. Scale bar 5 μm. Dashed line is the tip of the tail. C) Max projection of 200 nM Dynarrestin treated embryo. Scale bar is 5 μm. Dashed line is tip of tail. D) Max projection of 200 nM Ciliobrevin D treated embryo. Scale bar is 5 μm. Dashed line is tip of tail. E) Division of head neurons in untreated digested embryos. Arrowheads to show centrosomes Scale bar is 1 μm. F) Division of head neurons in 200 nM Dynarrestin treated digested embryos. Arrowheads to show centrosomes. Scale bar is 1 μm. G) Division of head neurons in 200 nM Ciliobrevin D treated digested embryos. Arrowheads to show centrosomes. Scale bar is 1 μm. H) Division of VNC cells in untreated digested embryos. Arrowheads to show centrosomes. Scale bar is 1 μm. I) Division of VNC cells in 200 nM Dynarrestin treated digested embryos. Arrowheads to show centrosomes. Scale bar is 1 μm. 0’ is the imaged immediately following treatment. J) Division of VNC cells in 200 nM Ciliobrevin D treated digested embryos. Arrowheads to show centrosomes. Scale bar is 1 μm. 0’ is the imaged immediately following treatment. K) Swarm plot showing time to complete cytokinesis in untreated, 200 nM Dynarrestin and 200 nM Ciliobrevin D treated embryos. L) Inset time course showing the progression of the centrosome along the dendrite in both untreated permeabilized embryos and permeabilized embryos treated with 100 nM dynarrestin. Arrowheads to denote centrosome and dashed lines to show dendrite. Scale bar is 1 μm. 0’ is the imaged immediately following treatment. M) Swarm plot showing time for centrosomes to leave the cell body and enter the dendrite in both untreated and 100 nM Dynarrestin permeabilized embryos. Embryos at greater than 14 minutes had centrosomes that were never observed leaving the cell body before the onset of twitching which occurs 14-30 minutes following the terminal division of the neuron.

Dynarrestin blocks the ability of dynein to bind to microtubules without affecting ATP hydrolysis^44^. We treated embryos with 200 nM Dynarrestin and observed the effect on cycling neuroblasts within the head and the progenitors of the ventral nerve cord labeled by the *cnd-1* reporter. Over the course of 30 minutes, we observed cells that show signs of initiating mitosis including the duplication of centrosomes, rounding of the cell body, and alignment of the centrosomes along the putative division axis, but were unable to divide within the normal period of time (5-10 minutes, based on observation of N=7 cells in 5 untreated animals). These cells persisted in this state for over 30 minutes (n=6 cells in 4 treated embryos) (Fig 4F/I). In both untreated embryos and undigested embryos treated with 200 nM dynarrestin neuroblasts within the head and the ventral nerve cord were able to undergo mitosis and continue development normally, such as tail elongation and dendritic extension of the head sensory neurons over the same time course (Fig 4B/C).

Another reported dynein inhibitor, Ciliobrevin D, acts via a distinct mechanism by inhibiting dynein’s ATPase domain, thus blocking its processivity^49^. We treated enzymatically digested embryos with 200 nM Ciliobrevin D, the same dose as Dynarrestin, and confirmed its impact on dynein activity by observing mitotic failures in neuroblasts of the head and ventral nerve cord (N=6 cells in 5 embryos). As with Dynarrestin, neuroblasts within the head and the ventral nerve cord arrested in mitosis while other developmental processes, including the elongation of the tail and the extension of sensory dendrites in the head, continued over the course of 30 minutes (Fig 4D/G/J). Having confirmed the efficacy of both drugs, we selected dynarrestin to use as our primary treatment as Ciliobrevin D exhibits a high level of autofluorescence as condensates appear in the medium over time which degrades fluorescence imaging.

We took advantage of the rapid onset of dynein inhibition based on the appearance of mitotically arrested cells within 5 minutes of treatment in our positive control experiments by precisely timing the addition of 100 nM Dynarrestin, just following the division of a single late-dividing sensory neuron on the ventral surface of the embryo’s head. The dose was titrated down to mitigate pleiotropic effects on the cell divisions. We then tracked the centrosome from drug addition up until the onset of embryonic twitching using an endogenously tagged allele of PCMD-1 (Fig 4K). In untreated embryos, the centrosome leaves the cell body and enters the dendrite in 3 ± 2 minutes (n=12 cells across 8 embryos). Upon Dynarrestin treatment, the centrosome ceases motion within the cell body and fails to migrate along the dendrite before the onset of twitching (n = 17/18 embryos, Fig 4L), suggesting that the activity of cytoplasmic dynein is essential for the timely export of the centrosome from the cell body following the terminal division of sensory neuroblasts.

## Discussion

The study of cell and developmental biology has long made use of an assorted toolkit of highly specific toxins, inhibitors, and fluorescent dyes to manipulate and illuminate molecular processes *in vivo*. The impermeability of the *C. elegans* eggshell has limited the adoption of these tools for the study of developmental processes, motivating several efforts to overcome these limitations. The discovery of *perm-1,* and its inhibition by RNAi to render the early embryo permeable to small molecules was a revolutionary step forward in this field and has been used to dissect diverse processes in early embryonic development. Despite its utility in studying events in early-stage embryos, the inability of *perm-1* RNAi embryos to develop beyond gastrulation has prevented its use in later stages of embryogenesis such as in the study of tissue morphogenesis or neuronal cell biology. Our determination that simplified cell culture medium support an exceptional level of embryo viability after eggshell removal through to larval development and adulthood provides a useful route forward for the introduction of these tools to the cell and developmental biologist lacking access to laser microbeams for targeted permeabilization, while reducing variability in preparation compared to the application of excess coverslip pressure. Given the high viability of *ex ovo* embryos cultured in this manner, permeabilization can take place well in advance of any developmental timepoint of interest, allowing for precisely timed drug addition for acute perturbations.

Cytoskeletal manipulations have been a major focus of prior strategies for eggshell permeabilization given the challenges involved in genetic manipulation of microtubule and actin networks. To combat these limitations the bulk of previous work has been limited to the first or first few cell divisions through a combination of both *perm-1* and genetic manipulations to the microtubule networks. We found that cultured *ex ovo* embryos are highly compatible with a range of commonly used toxins, often showing robust phenotypes at 0.1-1% of dosages reported in previous reports in *perm-1* RNAi, laser microbeam, and coverslip pressure permeabilized embryos. By combining fluorescently-tagged digested embryos and small molecule treatment, we are able to time the addition of those small molecules to exact developmental timepoints such as the start of ventral enclosure to target that event without disrupting normal development up until that point. Most notably, this temporal precision let us isolate a single step of neuronal morphogenesis: adding a dynein inhibitor within minutes of a sensory neuron’s terminal division showed that dynein activity is required to transport the centrosome into the nascent dendrite, a perturbation difficult to achieve genetically because dynein is essential and acts pleiotropically throughout development.

We have demonstrated that a wide range of drugs work effectively in these culture conditions at picomolar to nanomolar concentrations. Embryos cultured using this approach are not only viable but develop normally and are able to generate viable progeny, a marked improvement over *perm-1* RNAi, which compromises embryo viability beyond gastrulation, and without the potential acute mechanical stressors of laser microbeam and coverslip pressure induced permeabilization. Since eggshell digestion is scalable, large pools of *ex ovo* embryos can be easily prepared. The generation of *ex ovo* embryos and their culture will be a useful tool for cell and developmental biology, expanding the accessibility of molecular tools in the study of *C. elegans* embryonic development.

## Acknowledgements

This work was supported by NIH grant R35GM151199 to PKS. CAS was supported in part by NIH grant T32GM145388. Some strains were provided by the CGC, which is funded by NIH Office of Research Infrastructure Programs (P40OD010440). zyIs36 was a gift of Antonio Colavita. The RNAi library was provided by Patrick Allard. The authors would like to thank Jorge Torres and Margot Quinlan for the gift of reagents used in this study and valuable discussion.

## Methods

### C. elegans strains

**Table 1:** C. Elegans stains used in this study.

| Strain | Genotype | Source |
| --- | --- | --- |
| FT1197 | *xnIs449* [*lin-26::lifeAct::GFP* + *unc-119(+)*] | CGC |
| N2 |  | CGC |
| JH2054 | axIs1492 [pie-1p::GFP::tbb-2 ORF::tbb-2 3'utr + unc-119(+)] | CGC |
| TMD119 | pcmd-1(syb486[gfp::pcmd-1]) I | CGC |
| zyls36 | zyls36 [cnd-1p::PH::mCherry; myo-2p::mCherry] X | Shah & Tanner *et al.* 2017 50 |
| PKS19 | zyls36, unc-119(ed3) III; axIs1492. | This paper |
| PKS27 | zyls36, pcmd-1(syb486[gfp::pcmd-1]) I | This paper |

### Small molecules

**Table 2:**
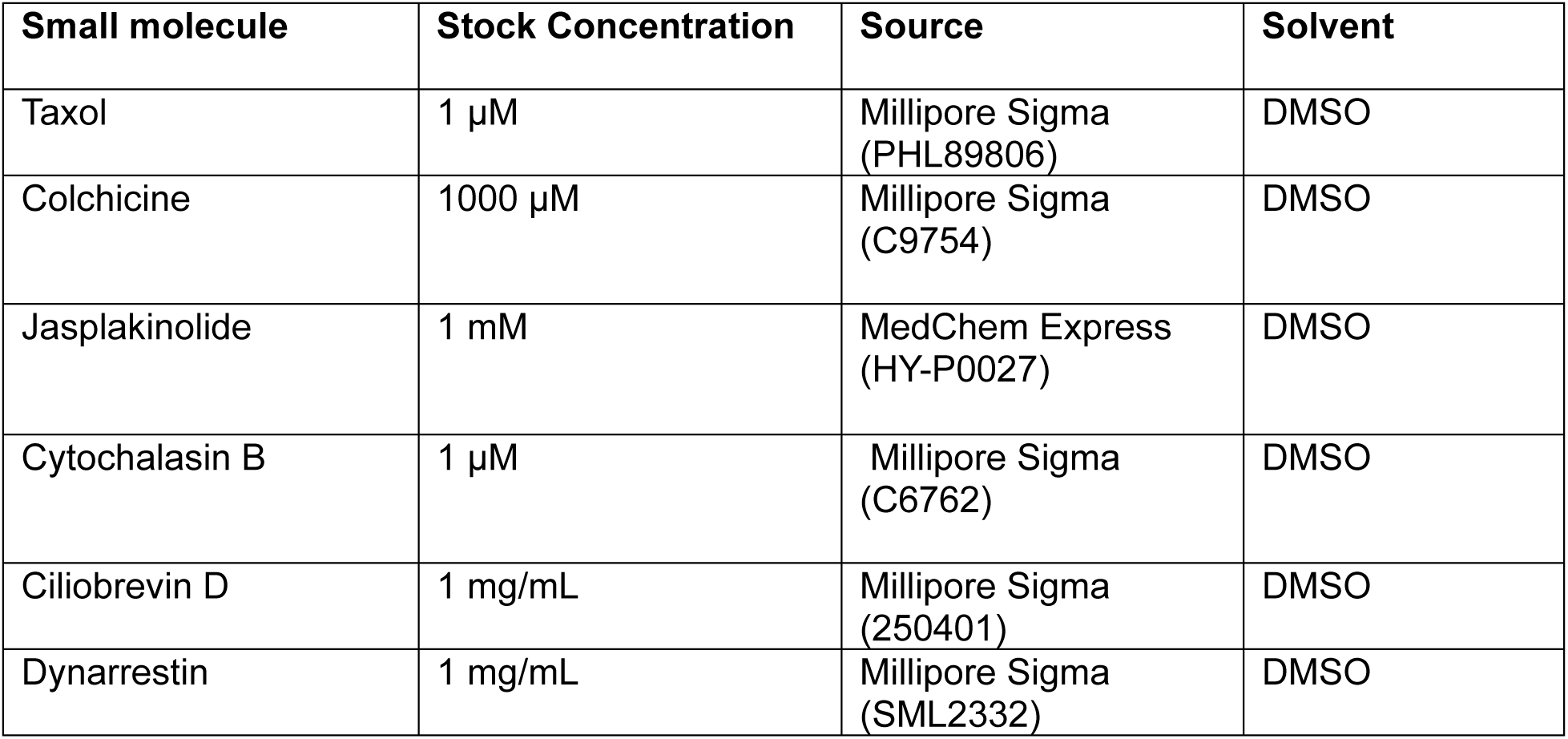
Small molecules used in this study.

Small molecule dilutions greater than 100X were done by serial dilution into L15 + 1.54% sucrose medium.

### Medium preparation

Supplemented medium was prepared by dissolving 7.7 g sucrose into a 500 mL bottle of commercial L-15 medium. The supplemented media was stored at -20 °C. At this temperature, the media did not appear to degrade over time (for a period of at least 6 months). After thawing, if the medium was used for long-term culture experiments (longer than approximately 2–3 hours), penicillin/streptomycin (100 units/mL final concentration) was added prior to use. Adding antibiotics prior to freezing resulted in precipitation that could interfere with imaging but did not appear to affect embryo viability.

### Embryo Collection

Adult worms were seeded onto an NGM medium plate for at least two days to obtain enough embryos. The plate containing embryos was washed off by pipetting 3 mL M9 buffer and collecting it into a 15 mL centrifuge tube. The buffer was repeatedly drawn up and ejected onto the plate to completely collect worms. If residual worms and embryos remained on the plate post-washing, an additional 1 mL of M9 was used. Enough M9 buffer was then added to the centrifuge tube containing worms and embryos to reach the 5 mL line. The contents were vortexed and centrifuged at 2,000 rpm (600 rcf) for 7 seconds to gently pellet the worms and embryos. The supernatant was aspirated down to the 1–2 mL line, taking care not to disturb the pellet. The pellet was washed again with 5 mL M9 solution three times as described above to remove bacteria.

5 mL of M9 buffer was added to the pellet. Then, 250 µL of 10-15% NaOCl (Millipore Sigma #425044) and 125 µL of 5M NaOH were added. The tube was vortexed in 30–60 second increments for 1–2 minutes. Embryos were pelleted by centrifuging at 2000 rpm for 6 seconds. The supernatant was aspirated, taking care not to disturb the pellet, and another round of the bleaching process was performed as described. After each vortexing step, the sample was checked under a dissection scope to confirm whether embryos were released from gravid adults and all worms fully digested to minimize debris carried over into subsequent steps. The bleached embryos were washed with 5 mL of the L-15 + sucrose + pen/strep medium three times to ensure complete removal of bleach.

See Figure 5

**Figure 5:**
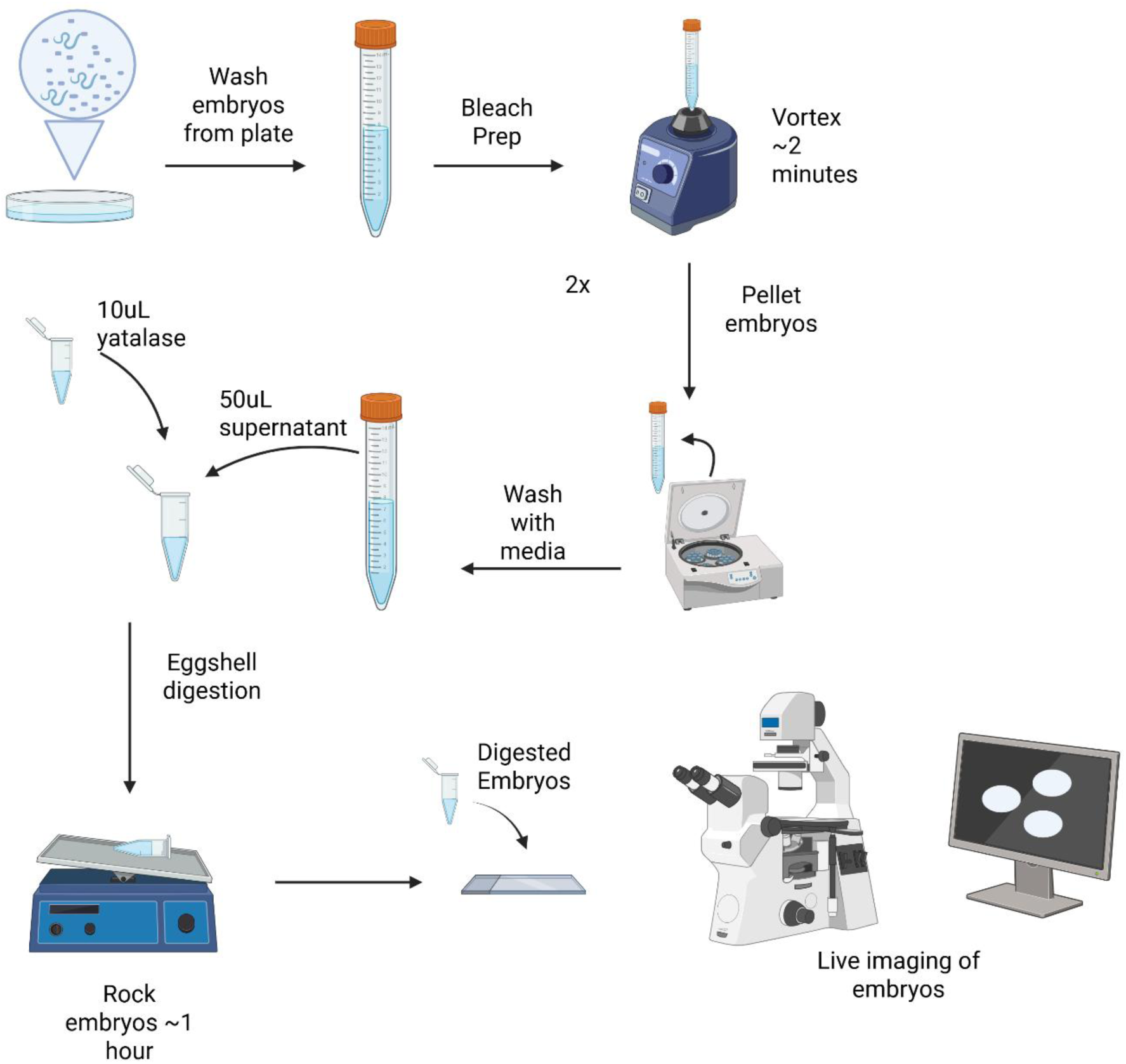
Schematic of embryo preparation. Created in BioRender. Stubbert, C. (2026) https://BioRender.com/p21cjln

### Eggshell digestion and embryo culture

After the final wash, embryos were pelleted again and excess medium was removed. 50 µL of the embryos remaining were transferred to a microcentrifuge tube, followed by 10 µL of 12.5 mg/mL Yatalase (Takara T017). Both single and strip microfuge PCR tubes have been used and it does not seem necessary to use nonstick tubes. If processing fewer than 100-200 embryos, a single 10 µL Yatalase aliquot was added and incubated on a rocking rack for 1 hour before proceeding to mounting. After this stage, we do not wash the remaining Yatalase from the solution to minimize handling and embryos can immediately be mounted for imaging and small molecule addition. Embryos can be left in the enzyme until they hatch with no effect on embryo viability.

If bulk processing large numbers of embryos (∼500 embryos), the suspension was incubated for 20 minutes while rocking after the first 10 µL Yatalase addition, and then left to stand for 10 minutes to allow the pellets to settle by gravity. Then, 10 µL of supernatant was carefully pipetted from the surface and examined under a microscope for the presence of embryos. A second 10 µL Yatalase aliquot was then added to the tube and mixed by pipetting up and down with a final volume of 50 µL. The sample was not vortexed at this stage, as partial digestion from the first Yatalase addition makes embryos fragile and prone to rupture. The embryos were incubated with the second Yatalase treatment for another 20 minutes on the rocking rack. Embryos are then mounted for imaging and small molecule addition from enzyme solution without washing away the enzyme. Embryo viability within the enzyme solution remains the same as with low embryo count digestion.

To verify the viability of *ex ovo* embryos, we transfer 10-20 embryos into 100 μL of fresh medium in a 35 mm petri dish, cover the droplet with mineral oil to prevent evaporation, and leave the embryos at room temperature overnight, scoring survival based on the appearance of swimming L1 larvae the following morning.

### RNAi

For RNAi, a clone of T01H3.4 from the Ahringer library was grown in 200 µL of LB + carbenicillin overnight at 37C^51^. 100 µL of bacterial culture was added to NGM plates with X M IPTG and left to dry overnight and induce expression. 20-40 L4 or young adult worms were added to each plate and left in a 20C incubator overnight before collecting permeabilized embryos.

For testing the viability of RNAi-permeabilized embryos, worms were transferred to a microwell slide in 50 µL of either 0.7x Egg Salts as originally described (1X Egg Salts: 118 mM NaCl, 40 mM KCl, 3.4 mM MgCl2, 3.4 mM CaCl2, 5 mM HEPES pH 7.4)^10^, or 1.54% sucrose in L-15 medium. 5 worms were cut using a scalpel and an eyelash was used to separate out 2-4 cell embryos. Those embryos were transferred to 24x 50 mm coverslips and mounted using ∼5 µL of 0.7x egg salts or 1.54% sucrose in L-15 medium with 20 *μ*m beads (42719). A 22 mm square coverslip was placed onto the droplet and the edges sealed with melted vaseline. Embryos were imaged for 4 hours on Olympus IX83 inverted microscope frame, a UPSAPO 60XS2 silicone oil immersion objective and a VisiTech iSIM super resolution multipoint scanning confocal microscope.

### Light microscopy

*Ex ovo* embryos are extremely fragile mechanically and care should be taken to minimize direct manipulation but are compatible with standard nematode embryo mounting practices such as bead-based compression mounting. Compression mounting with agar pads invariably results in the destruction of the embryos and should be avoided. We evaluated a handful of alternate sample mounting strategies in cases where the timing of drug introduction is desirable for temporal control of treatments for inverted high-resolution imaging.

Imaging was performed using either DIC alone for N2 or both DIC and fluorescence for all other lines using an Olympus IX83 inverted frame equipped with a UPLSAPO 60xs2 objective, a Visitech iSIM multipoint confocal scanner, ASI MX2000XYZ stage, and Hamamatsu Orca Fusion camera. The mCherry channel of zyls36 was acquired using 561 nm excitation and a 570 nm long-pass emission filter using 150 ms exposures and a laser power that was empirically tuned to not cause any qualitative developmental delays versus un-imaged control embryos and maintain a ∼100% hatch rate for imaged embryos. The GFP channel of both FT1197, PSK19 and PKS27 was acquired using 488 nm excitation and a 525/50 nm filter using 150 ms exposures and a laser power that limited photo-bleaching and maintained embryo viability. Embryos were imaged every 60 seconds with a 750 nm z spacing. DIC images were acquired with the Visitech scanner in brightfield bypass mode, a 50 ms camera exposure and the LED light source tuned to not generate any saturated pixels in the image. DIC illumination was generated using an Olympus UCD8 manual condenser equipped with a U525 oil immersion 1.4 NA top lens and a DICTHR tilt-shift slider. Images were acquired using micro-manager and cropped and converted to individual tiff volumes using Fiji. The DIC slider shear was adjusted empirically to approximate the contrast characteristics of images acquired on our Olympus microscope and images were acquired with a 50 ms exposure, 750 nm z-spacing, and illumination LED intensity adjusted to fill the sensor dynamic range without saturating pixels.

### Bead mounting for imaging with inverted microscopes

*Ex ovo* embryos tolerate compressed mounting using methods previously optimized for imaging during long-term time lapse for cell lineage tracing^52^. Briefly, a suspension of 20 µm diameter polystyrene beads (Thermo Scientific 09-980-133) is prepared in L-15 medium with sucrose and antibiotics. This preparation is diluted empirically to achieve a final concentration of 100-200 beads per µL. For imaging, *ex ovo* embryos are transferred into a 2 µL drop of bead suspension on a #1.5 coverslip by capillary micropipette. A smaller coverslip is placed on top and secured in place by painting with melted petroleum jelly without moving the coverslip after placement to avoid damaging the embryos. Imaging post-twitching embryos in this format is made challenging as the lack of a rigid eggshell allows *ex ovo* embryos to generate traction against the coverslip surfaces and migrate slowly. This mounting set up does not easily allow for the timed addition of small molecules following imaging unlike the open mounting and was used for the fluorescent dye imaging, microtubules dynamics and actin dynamics imaging.

All imaging was performed using an Olympus IX83 inverted microscope frame, a UPSAPO 60xs2 silicone oil immersion objective and a VisiTech iSIM super resolution multipoint scanning confocal microscope.

### Open mounting for accessibility for drug delivery

While bead-compressed mounting is advantageous for numerous applications, the closed nature of the preparation complicates acute addition of small molecules. For the timed treatment of embryos with the dynein inhibitors Dynarrestin and Ciliobrevin D, we used open-chambered slides (ibidi 80821) to mount embryos. Mounting embryos in 200 μL of medium and diluting additives to 100✕ stock concentrations to reduce the volume of the added drug minimized disruption to the embryo positions during time lapse imaging, allowing for precisely timed drug addition.

## Notes

### Competing Interest Statement

The authors have declared no competing interest.

### Summary of Updates

Additional results added for dynein inhibition, significant revision of the main text

